# Click-linking: a cell-compatible protein crosslinking method based on click chemistry

**DOI:** 10.1101/2024.12.23.630117

**Authors:** Bruno C. Amaral, Andrew R.M. Michael, Nicholas I. Brodie, D. Alex Crowder, Kristen H. Eiriksson, David C. Schriemer

## Abstract

Crosslinking mass spectrometry (XL-MS) has the potential to map the human interactome at high resolution and with high fidelity, replacing indirect, error prone sampling methods such as affinity pulldown MS. However, the sampling depth of XL-MS remains stubbornly low. We present a novel crosslinking strategy that splits the crosslinking reaction into two sequential and orthogonal coupling events. The method involves pre-stabilizing the spatial proteome with a fixation protocol inspired by immunofluorescence imaging, followed by a stepwise process that begins with extensively labeling surface-accessible lysines in the cell with N-hydroxysuccinimide (NHS)-modified click reagents. We show that a subsequent copper-catalyzed azide-alkyne cycloaddition (CuAAC) reaction of the installed precursors generates crosslinks at levels approaching 30% of the total signal, as demonstrated by a subtractive approach. The method generates no detectable side reactions or obvious distortions of the spatial proteome. Protein-protein interactions (PPIs) are detected at levels approximately 20 times higher than a conventional DSS-based method, outperforming even enrichable crosslinkers.

## Introduction

Protein-protein interactions (PPIs) are a major organizational element in the cell. Mapping them as comprehensively as possible is necessary to understand the mechanisms behind every cellular process^1^. Mapping can be performed at different resolutions and scales. Structural techniques such as X-ray crystallography and cryoelectron microscopy provide the highest resolution and deliver the greatest mechanistic information about PPIs, but they have limited utility as a discovery tool and cannot be easily implemented at the cellular level. Techniques such as affinity pulldown mass spectrometry (AP-MS)^2^ and Bio-ID proximity labeling^3^ can be used for discovery, along with modern variants of yeast two-hybrid assays^4,5^, but the structural resolution of these techniques is limited, and PPIs are only inferred from the resulting data. Most of these methods are labor intensive when applied at the cellular scale, and they struggle to map and monitor dynamic interactions across time. We still need a robust technique that directly maps PPIs in cells, and one that is easy to use.

Crosslinking mass spectrometry (XL-MS) could be that superior method. It provides direct evidence of an interaction by defining a PPI based on a measured distance between two covalently bound proteins, through the insertion and detection of a bifunctional reagent of known length^6^. *In situ* crosslinking experiments have been moderately successful in PPI mapping^7,8^.

PPIs have been detected that agree with existing interaction knowledgebases and novel interactions are beginning to emerge, at least among proteins of higher abundance. Recent studies have even demonstrated that *in situ* crosslink measurements can support AI-driven structure modeling^9,10^, raising the possibility of determining protein structures in the cell. Unfortunately, *in situ* applications of XL-MS do not deeply sample the interactome. Some success has been achieved, but these experiments are a significant undertaking in terms of time and cost, while still only producing modest numbers of PPIs^11–14^. The crosslinking step seems straightforward – simply bathing live cells with a solution of the crosslinking reagent – but the sample workup necessitates large amounts of lysate and elaborate detection methods. New approaches are needed.

There are at least two experimental issues to address. First, there is some uncertainty in the accuracy of PPIs identified via XL-MS, as kinetic trapping could generate non-native interactions. Second, the yields of crosslinking reactions are poor^15^. In a typical cell-based method, live cells are bathed with reagent and left to react for a long period of time (up to an hour), as a concession to the low membrane permeability of most reagents and the need to integrate reaction products to detectable levels. These conditions are not conducive to preserving cellular structures, nor to a error-free sampling of interactome dynamics. We recently showed that a pre-stabilized cell retains structure during subsequent crosslinking reactions, and reaction yields are improved at the same time^16^. This stabilization is achieved with high-concentration formaldehyde in a method inspired by standard immunofluorescence (IF) imaging. We simply replace the fluorescent stain post-fixation with MS-compatible crosslinkers. Prestabilization allows *in situ* XL-MS to inherit the same level of confidence in PPI detection as IF imaging. Any type of crosslinker can be introduced post-fixation, even membrane-impermeable ones. We call this method formaldehyde-enhanced *in situ* crosslinking MS (FIX-MS).

FIX-MS does improve performance, but crosslinking reaction yields are still not as high as they need to be. Yields hover around hundreds of PPIs even after extensive sample separation and enrichment, at least when beginning with low milligram amounts of protein^16^. Interactome coverage remains a small fraction of the estimated 650,000 pairwise protein interactions present in human cells^17^. We were interested to see if reaction yields could be improved and there are reasons to think that it is possible. Typical NHS-based amine-targeting reagents, by far the most common, can partially hydrolyze during the crosslinking reaction and block otherwise linkable lysines. These reagents may also interact with other biomolecules, and even self-compete^18^.

Thus, monolinks are the dominant product^15^. This problem occurs because crosslinking methods use one molecule with two identical reaction groups, in a concerted reaction.

Uncoupling the stabilization of the cell from the insertion of crosslinks provides an opportunity to reinvent *in situ* crosslinking. We propose to separate crosslinking into two orthogonal reactions (**Fig. 1A**). For example, a first coupling event could support the labeling of surface-accessible lysines, followed by a washing step to remove the remaining reactants and at least some reaction byproducts (*i.e.*, hydrolyzed reagents). Initiation of the second coupling, which creates the crosslink, could then occur in an environment where insertion yields can be measured and controlled, and any competition is only between inter and intra-protein linkage events. Here, we investigate this concept using click chemistry as the second reaction and test its merits. We demonstrate that this “click-linking” approach is compatible with the FIX-MS protocol and highlight the potential of the method for deep PPI sampling.

**Figure 1.**
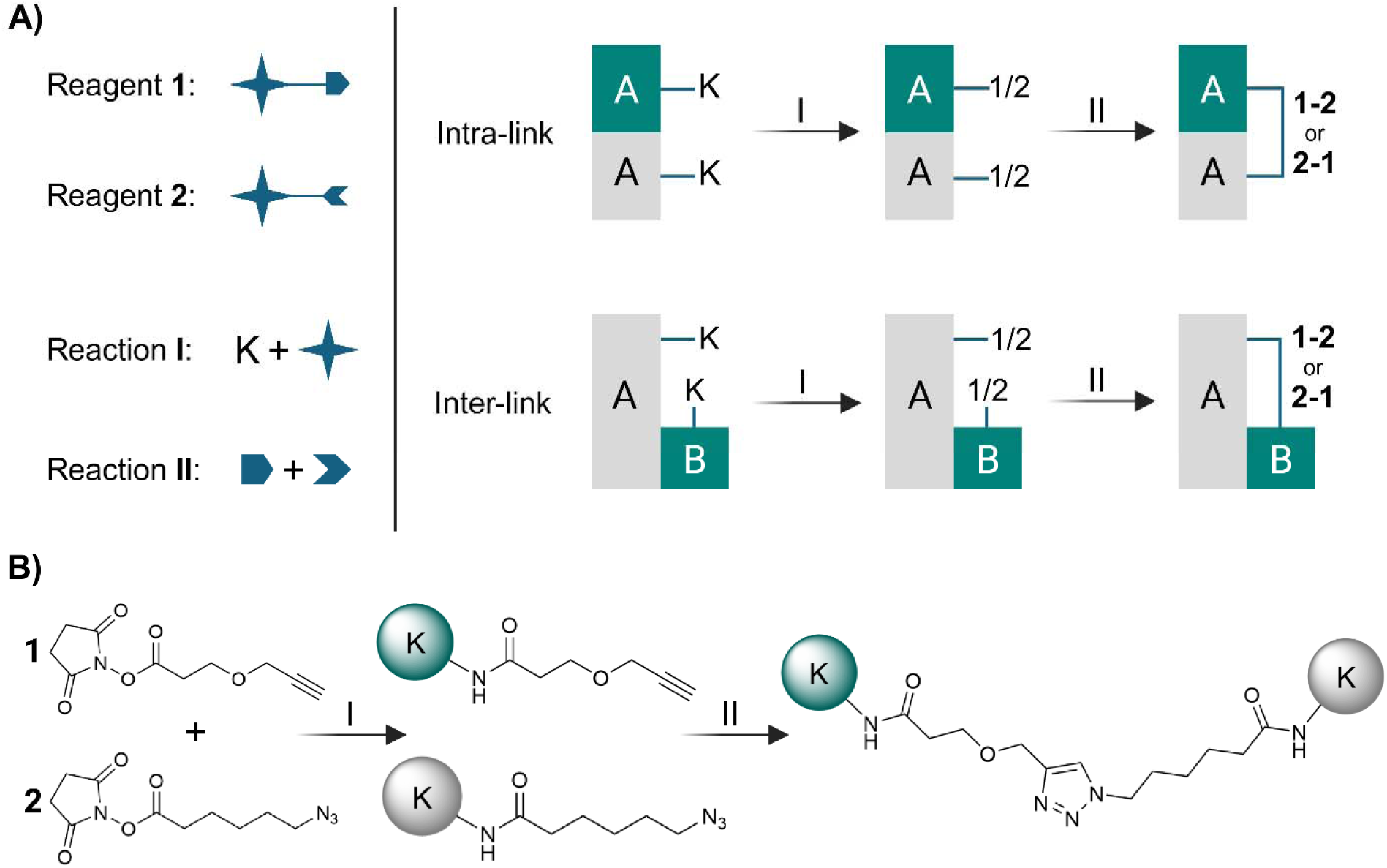
Deconstructing the protein crosslinking reaction. (A) Conceptual approach to deconstructing crosslinking into two orthogonal reactions. To illustrate, the first reaction can target amines, where both reagents **1** and **2** compete for available lysines. When admixed in a 1:1 molar ratio and applied to a stabilized cell, the reagents can label all accessible lysines with controllable yields. Each amine will possess a 1:1 ratio of the products of **1** and **2**. In the two examples provided, an intralink or an interlink will form after initiation of the orthogonal reaction, to a maximum of 50% of the bioconjugation yield (assuming linkage to only one proximal amine is possible). Asymmetry in the second reaction would lead to two reaction products (same mass, different orientation). (B) Click-linking reaction scheme used in this work. 2,5-dioxopyrrolidin-1-yl 3-(prop-2-ynyloxy)propanoate (**1)** and (2,5-dioxopyrrolidin-1-yl) 6-azidohexanoate (**2)** contain NHS esters for a standard lysine bioconjugation (reaction I). Reaction II is based upon copper-catalyzed azide-alkyne cycloaddition (CuAAC), using Cu(I) and BTTES.

## Results and Discussion

### Deconstructing the crosslinking reaction

Separating crosslinking into two sequential reactions first requires the selection of orthogonal coupling chemistry (**Fig. 1B**). We chose N-hydroxysuccinimide (NHS) esters for the first step, to selectively bioconjugate free amines in lysines and protein N-termini. Retaining a high degree of selectivity on the primary coupling will facilitate the detection of any crosslinks that we create, as database searches can be targeted to a reduced set of residues. While other chemistries are certainly possible (*e.g.*, coupling to carboxylic acids), NHS esters are very reactive and a wide range of coupled, secondary reactive groups are readily available. We selected reagents that enable copper(I) catalyzed azide-alkyne cycloaddition (CuAAC) reactions^19^. This click reaction generates 1,2,3-triazoles that are stable in solution and during MS analysis. The reaction is highly selective and its precursors are chemically compatible with NHS esters. The two precursors, when clicked, would form a reagent with a total spacer length of approximately 18Å, only slightly longer than most conventional crosslinkers. A catalyzed reaction is essential. The precursors will remain inert during bioconjugation in the absence of the catalysts, and only active in the subsequent step when the catalysts are added. Cu(I), together with activating ligands, increases the reaction rate by approximately 7 orders of magnitude^19,20^. Cu(I) is cytotoxic to live cells, but this is a non-issue as cells are fixed with formaldehyde at the start of the procedure.

The two precursors can be installed simultaneously in a 1:1 molar ratio. Barring any insertion bias, every free amine would possess an equimolar amount of each click precursor, and the crosslinking yield would be no worse than 50% of the combined insertion levels. For a single crosslinkable pair of lysines, azide would not couple to azide, nor alkyne to alkyne, but if the number of nearby linkable lysines increases, the crosslinking yield can approach 100%. For example, six linkable lysines proximal to a labeled lysine site would generate a 98.4% maximal crosslinking yield at the site.

### The reaction – step one

We fixed and permeabilized human A549 cells using the conventional FIX-MS approach, and then added 2,5-dioxopyrrolidin-1-yl 3-(prop-2- ynyloxy)propanoate (**1**) and (2,5-dioxopyrrolidin-1-yl) 6-azidohexanoate (**2**) in equimolar amounts. A combined concentration of 2 mM and a 60 min reaction time generated a high level of labeling. Using peptide intensity as a measure, 36% of the total signal arises from labeled peptides, or 61% when counting only lysine-containing peptides. This level can be boosted with repeat applications of the reagents to improve depth of labeling (**Fig. 2A**). We used the “1x” application of reagents for the rest of the reaction assessment.

**Figure 2.**
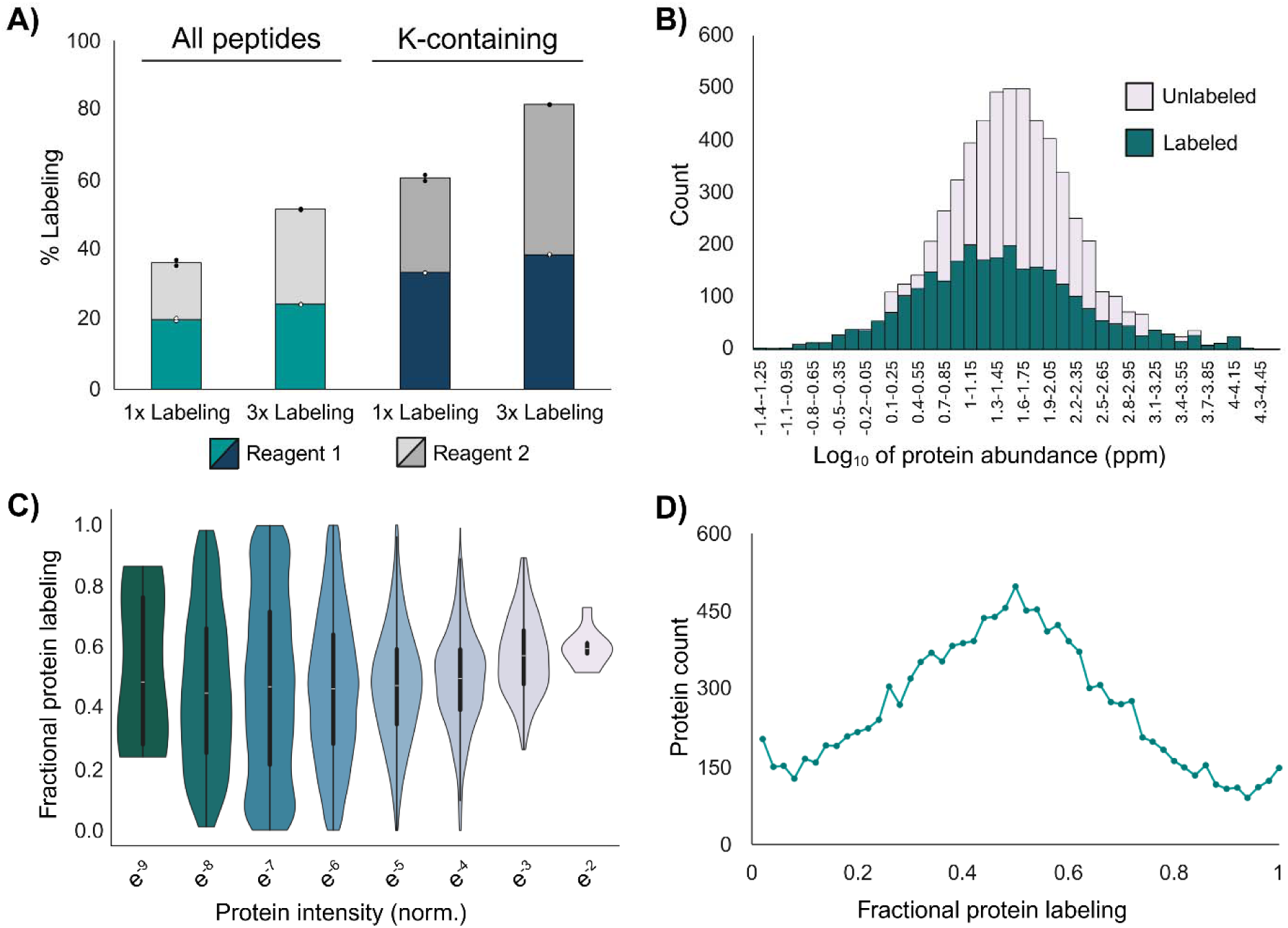
Properties of the initial NHS-based bioconjugation reaction. (A) Labeling levels achieved in a single application of a 1:1 molar ratio of reagents **1** and **2**, calculated by using all detected peptides as well as only the lysine-containing peptides. Labeling is expressed as fraction of total labeled peptide intensities over all peptide intensities for a given peptide class. Data showing a single application of reagent mix (1x) and three sequential applications of the reagent mix (3x). (B) Proteome-level distribution of labeling for the 1x application, relative to an unlabeled proteome analyzed in the same manner. A protein was counted as labeled if one or more labeled lysine-containing peptides was detected. Protein abundance estimates from the PaxDB. (C) Data as in B, with violin plot showing the distribution of fractional protein labeling (total labeled lysine-containing peptide intensity over total peptide intensity) as a function of protein abundance (estimated from peptide intensities normalized to total ion intensity). (D) Data as in B, showing a distribution of per-protein fractional labeling levels.

Approximately 90% of the labels that we install are on surface-exposed lysines, estimated by probing the reagent-accessible surface area (RASA) of several thousand AlphaFold protein models (**Supplementary Figure 1**). The bioconjugation yields for the two reagents are nearly equal. Minor deviations likely reflect dispensing errors, a modest insertion bias, and/or unequal identification probabilities for peptides that contain different modifications. The labeling depth is broadly reflective of the composition of the proteome. We appear to sample across the dynamic range of protein expression (**Fig. 2B**). The modified peptides are shifted towards the lower abundance elements of the proteome, but the shift is modest and likely arises due to altered digestion patterns. Labeled peptides render lysines resistant to trypsin and could shift the detection properties of the proteome. The labeling yields for individual proteins are variable but the average labeling per protein is generally independent of protein abundance (**Fig. 2C,D**).

We noted a strong bias against azide-modified peptides initially, but we speculated that these peptides experienced a greater level of loss during sample workup. Thus, we capped the bioconjugated azides before sample workup, using 11,12-didehydro-γ-oxodibenz[b,f]azocine- 5(6H)-butanoic acid (DBCO-acid). DBCO-acid is a very reactive strained alkyne that reacts with azides in a spontaneous strain-promoted alkyne-azide cycloaddition (SPAAC) reaction^21^. This capping returned detectability to the azides. No additional products of the bioconjugation reactions were found in an open-modification search, confirming the high selectivity of reaction.

### The reaction – step two

We then explored the effect of catalyst composition and reaction time on the ability to achieve an efficient click reaction, initially using cell lysates and a “half reaction”. That is, we installed **1** alone into *E. coli* and A549 cell lysates, and then added 2-[2-(2- azidoethoxy)ethoxy]-ethanol as a capping reagent. These monovalent modifications are much easier to detect with current proteomics methods than crosslinks. Classical CuAAC ligands such as THPTA, while effective, are strongly hydrophobic and they interfere with LC-MS analysis (not shown). Additionally, they require high concentrations of organic solvent in the reaction and could possibly induce cellular distortion, even after fixation. Thus, we used a newer generation ligand, 3-(4-((bis((1-(*tert*-butyl)-1*H*-1,2,3-triazol-4-yl)methyl)amino)methyl)-1*H*-1,2,3-triazol-1- yl)propane-1-sulfonic acid (BTTES), which is water soluble^23^. Aside from this one change, standard CuAAC reaction conditions proved highly effective, leading to a near-quantitative conversion of the propargyl group to the triazole in 15 min using a 2:1 ratio of ligand to Cu(I) (**Supplementary Figure 2**). The reaction time can be even shorter, but we elected to use 15 min as a default for all subsequent experiments. We did not observe any side products in the proteome, consistent with the bioinert nature of the CuAAC reaction. We note that a minor thiotriazole product could form with proximal cysteines, but it would be difficult to detect^24^.

### Click-linking – in situ click reaction of installed precursors

Using the reaction as described above, we then explored the effectiveness of the entire click-linking process, from fixation to installation of the click precursors, and then to the final cycloaddition reaction to create the crosslinks (**Fig. 3**). We chose a subtractive approach to evaluate performance. That is, we monitored the drop in the level of detectable click precursors because, as noted above, monovalent modifications are much easier to enumerate than crosslinked peptides. We reasoned that quantifying the drop in monovalent modifications would support a more accurate first estimate of crosslink success. The drop was substantial and reproducible (**Fig. 4**). In the 1x labeling experiment, the alkyne-modified peptides dropped by 80% and azide-modified peptides by 89%.

**Figure 3.**
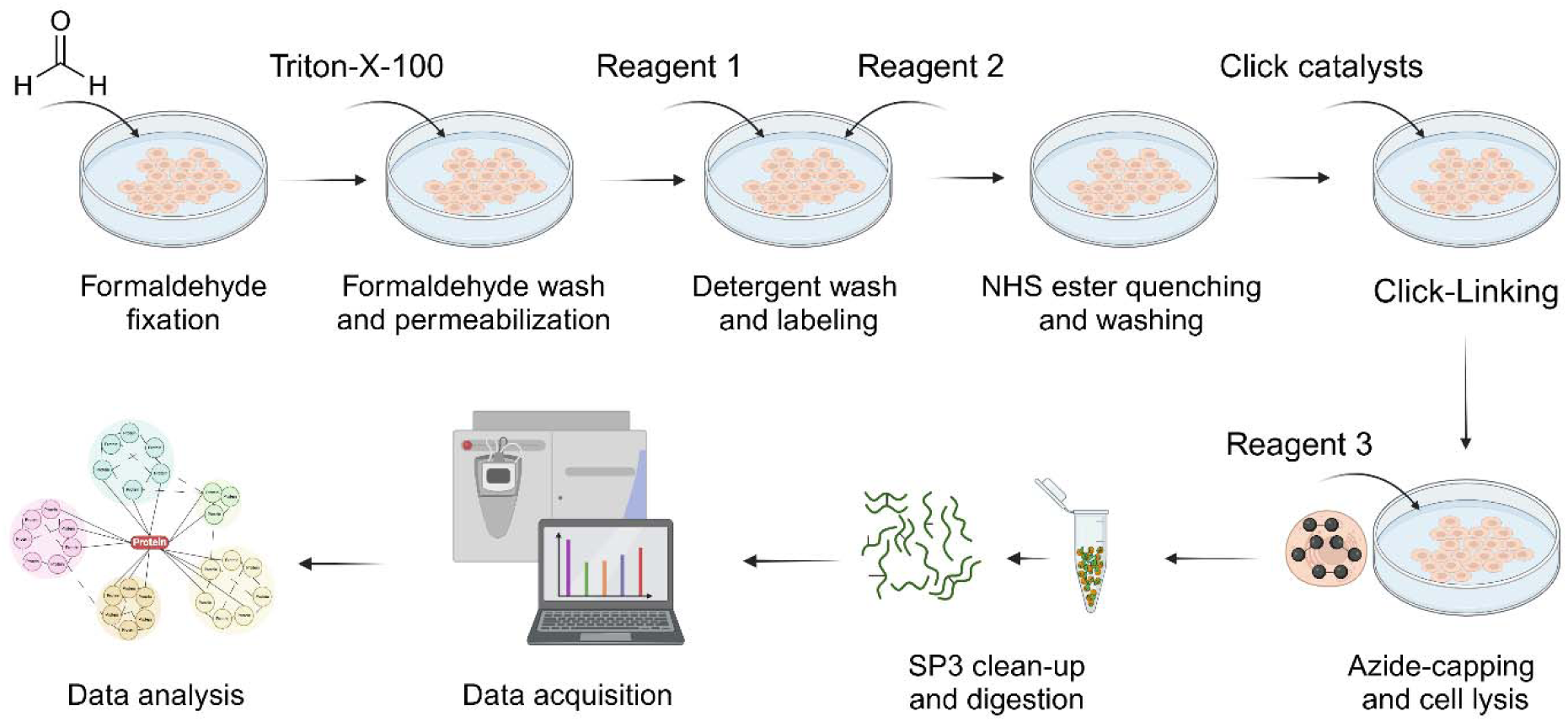
Schematic of the multistep click-linking workflow.

**Figure 4.**
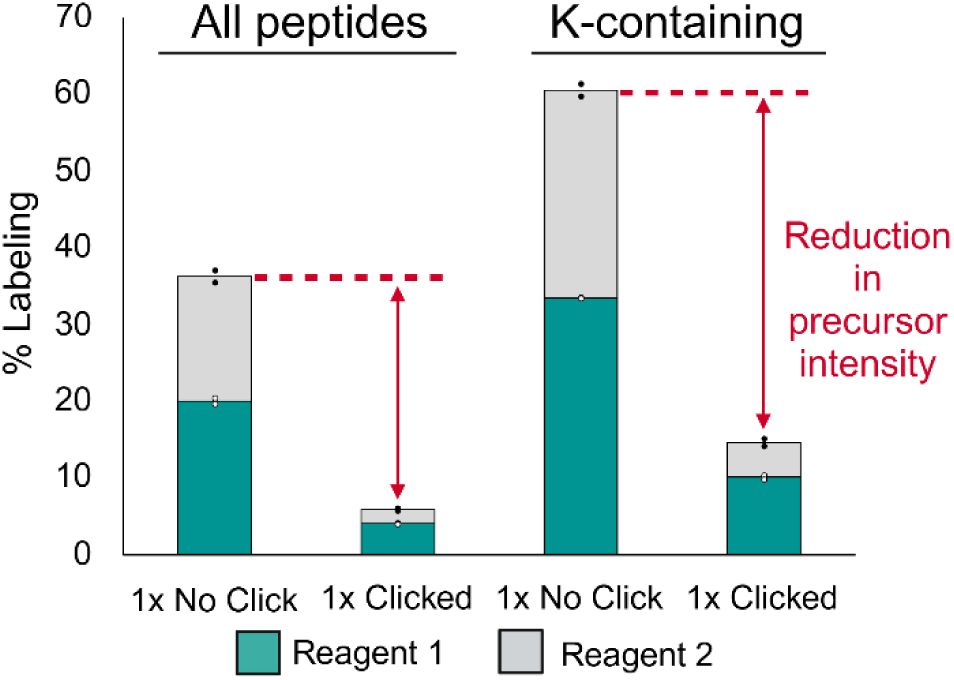
Effect of the click reaction on the abundance of detectable click precursor peptides. The reduction in detectable labeled peptides is expressed as a fraction of the total detectable signal and stated for all peptides and those only containing lysines.

We then considered what other biomolecules might possibly compete with the formation of protein crosslinks. We do not expect DNA or RNA to contribute crosslinkable sites, as they are not reactive to NHS esters. While it is possible that amine-containing metabolites could compete with proteins, the clink-linking process should deplete the cell of metabolites, as multiple washing steps are incorporated post-fixation. To test this, we extracted the metabolome from fixed and permeabilized cells and performed a quantitative metabolomics analysis of amine-reactive metabolites. We used live cells as a control, to represent the state of a conventional *in situ* crosslinking reaction. The metabolite levels are very high in the control, but the click-linking workflow reduces the amine-reactive pool of metabolites to nearly undetectable levels (**Supplementary Figure 3**). Interestingly, side reactions with metabolites are not seen in the conventional method. There are hundreds to thousands of amine-containing metabolites in a cell, thus signal splitting would likely render them undetectable in typical experiments. Taken together, the reduction in click-link precursors that we observe in **Fig. 4** is solely due to the production of intraprotein and interprotein crosslinks, suggesting a very high-yielding reaction of approximately 30% for the 1x reaction, considerably higher than the 0.1% estimate from previous studies of conventional crosslinking methods^15^.

### Click-linking does not distort cellular ultrastructure

We previously showed that the spatial proteome remained stable over multiple successive applications of crosslinker during FIX-MS^16^. However, the process of click-linking is slightly different than FIX-MS. If reversible formaldehyde crosslinks between amines form the basis for proteome stabilization^25^, then it is possible that an extended treatment with the NHS-based click precursors could outcompete the formaldehyde crosslinks and destabilize the cell. To evaluate this, we fixed and permeabilized A549 cells as usual, and then treated them with the 1:1 molar ratio of **1** and **2** (total concentration of 2 mM) for a total of 60 min. We then imaged the cells using CF647-phalloidin staining to observe the effect on the actin cytoskeleton, a structurally sensitive state that is coupled with many cellular processes^26^. The cells remained fixed and stable relative to an unlabeled control (**Fig. 5A**). Interestingly, reversible lysine-formaldehyde crosslinks do not appear to significantly contribute to the fixation process that we use in click-linking.

**Figure 5.**
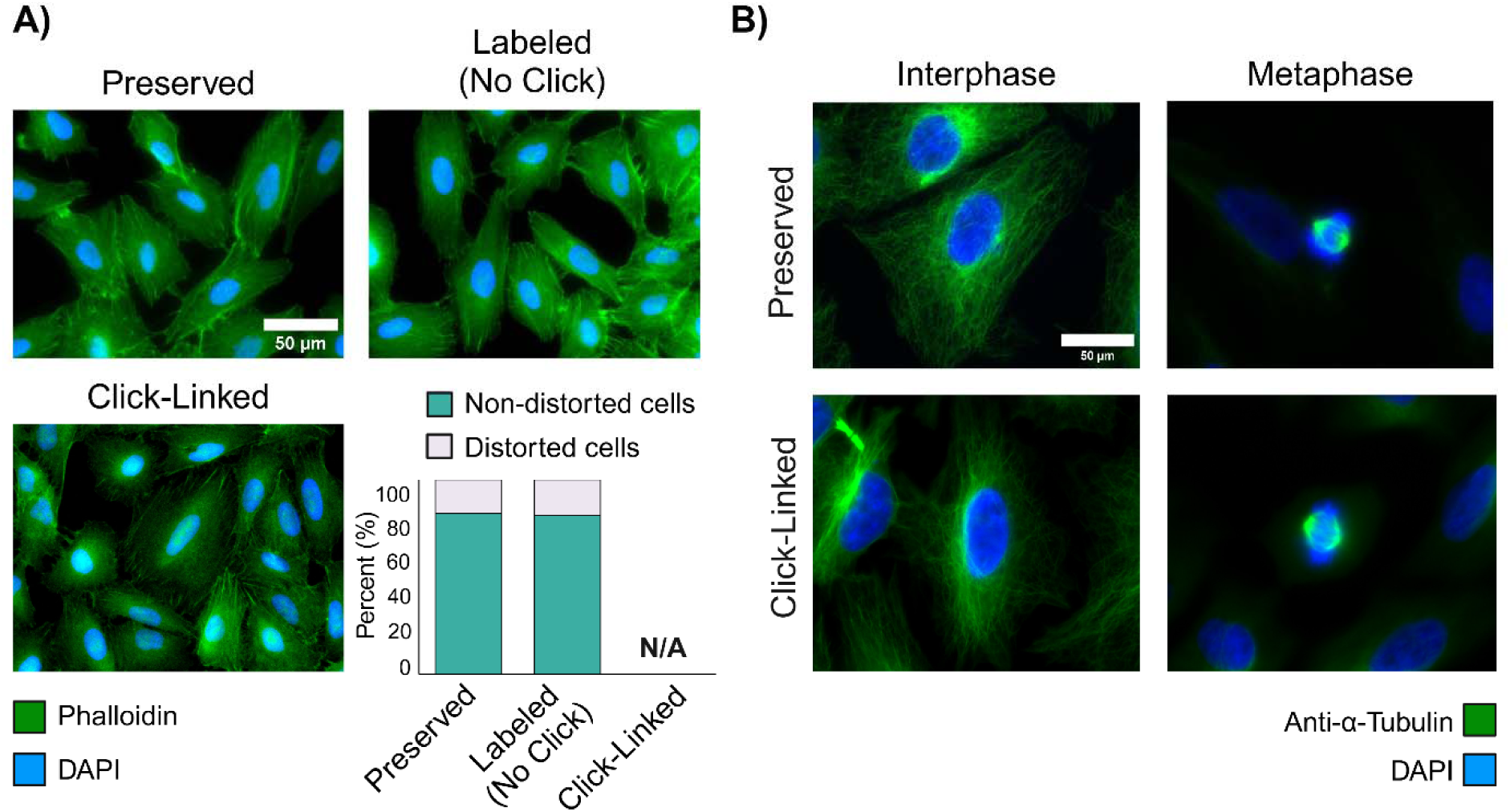
Click-linking does not negatively affect the ultrastructure of the cell. (A) A549 cells imaged using FITC-phalloidin (filamentous actin, green) and DAPI (nucleus, blue), after a standard cellular fixation protocol (top left), insertion of click precursors **1** and **2** to a 50% labeling level (top right), and after clicking and an EDTA wash (bottom left). Enumeration of cellular integrity in bottom right, based on defined metrics for integrity (see methods). (B) A549 cells imaged using an anti-α-tubulin antibody (green) and DAPI, after a standard cellular fixation protocol showing both a cell in interphase (top left) and mitotic cell in metaphase (top right). Cells imaged after the full click-linking protocol, also showing a cell interphase (bottom left) and a mitotic cell in metaphase (bottom right).

We then explored the effect of the click reaction. However, we were not able to sensitively image the actin cytoskeleton after introducing the click catalysts, as Cu(I) appears to interfere with phalloidin staining^27,28^. We treated the cells with EDTA to chelate and wash away soluble copper after click-linking. This process restored some of the signal, sufficient to indicate that the ultrastructure is preserved, but it was difficult to accurately quantify cell distortion (**Fig. 5A**). To confirm stability we stained for microtubules, another dynamic element of the cytoskeleton, after confirming that staining levels are unaffected by the reagents used in the click-linking process (not shown). There is no obvious change in microtubule structure upon click-linking. Even the fragile machinery of the mitotic spindle in metaphase is preserved (**Fig. 5B**), supporting our observation that the spatial proteome remains stabilized during click-linking.

### Click-linking outperforms standard crosslinking and FIX-MS

We then processed the “1x” click-linked A549 cells for XL-MS experiments, to compare crosslinking performance with two other well-known NHS-based crosslinking reagents, DSS and PhoX. We chose non-cleavable reagents so that the database search routines and parameters would be comparable.

DSS is a conventional non-enrichable reagent while PhoX is a highly-enrichable reagent^29^. Both were implemented using the FIX-MS process to increase insertion yields, and we also included a conventional live-cell treatment with DSS for comparison (data retrieved from an earlier study^16^). Click-linking generates CSMs and unique crosslinks that are slightly higher in number than DSS installed by FIX-MS, but the distribution of reaction products is quite different (**Fig. 6A,B**). Click-linking generates a much larger ratio of structurally informative crosslinks. Fully 60% of the crosslinks return structurally useful distance measurements (inter and intraprotein crosslinks), compared to 20% for DSS and 31% for PhoX. Indeed, while click-linking generates 60% less crosslinks than PhoX, the number of PPIs is greater (**Fig. 6C**). This is a remarkable result given that click-linking does not incorporate a strong enrichment mechanism like PhoX.

**Figure 6.**
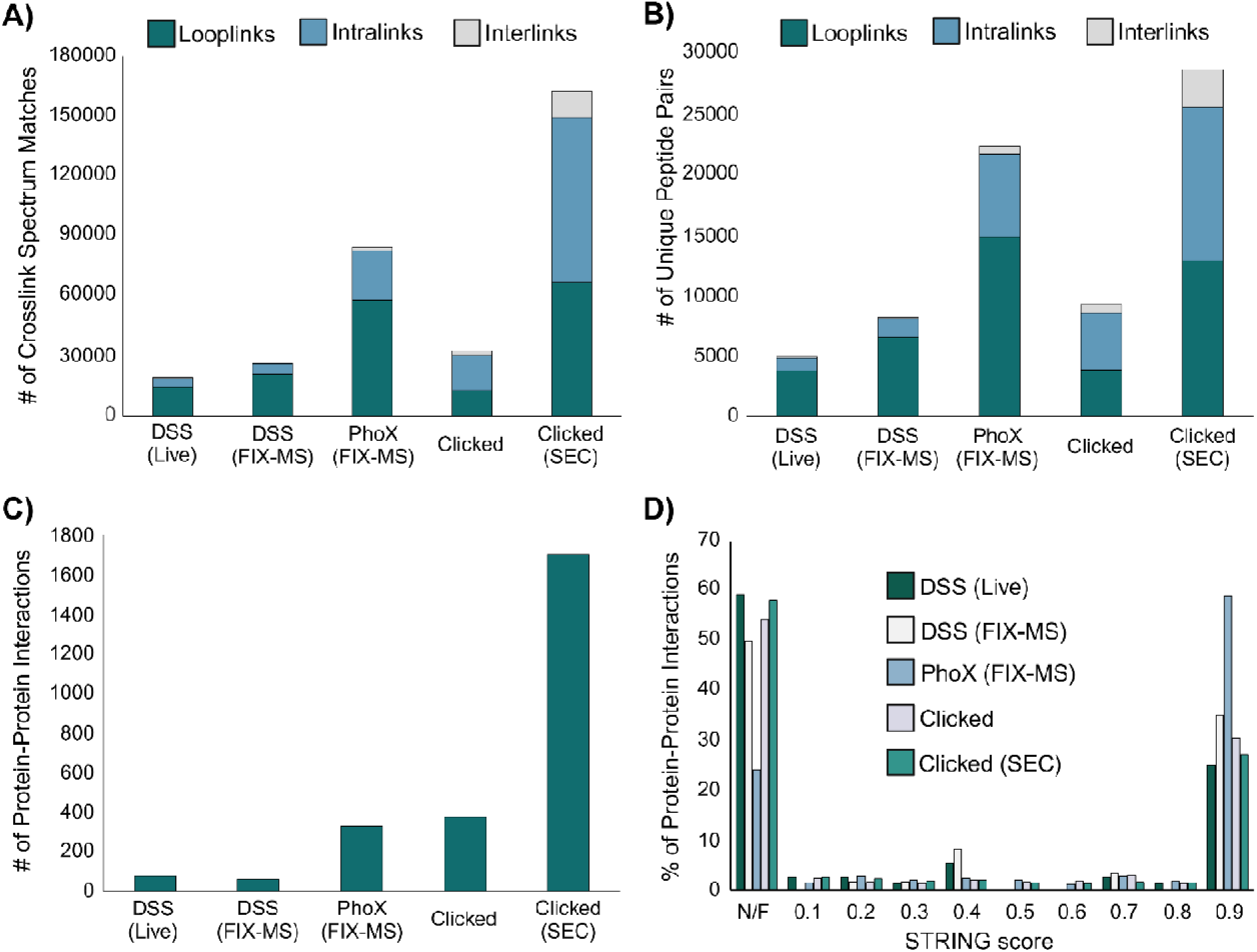
Comparing click-linking to alternative *in situ* XL protocols. (A) Number of crosslinked spectrum matches (CSMs) for a range of methods: live cells crosslinked in the conventional fashion with DSS, DSS and PhoX applied in a FIX-MS protocol, and the click-linking protocol. Click-linking results are also shown with an extra dimension of separation applied (size exclusion chromatography, SEC). (B) as in A, bur for unique crosslinked peptides. (C) As in A but displaying the number of unique PPIs detected. (D) Cross-referencing the PPIs detected by all methods against the STRING database.

There are at least two reasons for an improvement in PPI sampling relative to looplinks. First, the click-linked product is longer than DSS (11.4 Å) and PhoX (5 Å), which may generate more intra and inter-protein crosslink reactions. Second, the regioselective click reaction involves transition states that may be sterically unfavored when two click-linking precursors are installed close to each other on the same protein^30^. Additional reagent synthesis will help uncover if this observation is unique to click-linking. As the complexity of the reaction products is very high in XL-MS (and these products all compete for finite MS detection time) any tilt towards a higher percentage of interprotein crosslinks should improve PPI detection.

### In-depth analysis of the in situ click-linking reaction

These click-link precursors are not yet enrichable, thus linear free peptides still comprise the largest fraction of identifications from a typical experiment, even for click-linking (86%, **Supplementary Table 1**). We added an additional fractionation step to the workflow as partial enrichment (although this extra dimension is not as effective as the IMAC enrichment used for PhoX) and then measured the number of PPIs with an equally modest sample input (∼10 mg of protein). Size-exclusion chromatography (SEC) was used to fractionate the protein extract from a typical click-linking experiment. We generated five SEC fractions and then separated each fraction into additional fractions using high-pH reversed phase chromatography, prior to LC-MS/MS (see methods). This effort identified 1700 PPIs, five times greater than PhoX, and the advantageous distribution of reaction products was preserved (**Fig. 6A-C**).

We evaluated these 1700 PPIs against the STRING database and noted that, while a large fraction was corroborated with high STRING scores, an even larger fraction was not found in the database. This ratio is reproducible and generally independent of how the datasets were collected (**Fig. 6D**). Interestingly, the fraction of PPIs not found in STRING is higher for click-linking than for PhoX. There could be several reasons for this higher ratio of unknown PPIs. They may be false hits arising from a high error rate using our chosen database search tool (pLink2).

Additionally, if protein movement is not fully restricted upon fixation, indirect interactors could be linked. However, this should also negatively affect the PhoX-based method, but it does not appear to as the ratio is lower.

But is also possible that we are sampling novel PPIs, as the STRING database is not comprehensive. It is impractical to validate all 990 entries in this class, thus we sought to globally estimate boundaries on the error rate. We analyzed the gene ontology (GO) for the proteins comprising each PPI, reasoning that crosslinked proteins would derive from a common cellular compartment. Localized diffusion should not scramble this basic GO term. An analysis of all hits with a STRING score greater than zero showed that 667/710 (94%) of the PPIs were comprised of proteins drawing from the same cellular compartment, suggesting its utility as a filter. A random sampling of proteins in the proteome returns a 21% likelihood that PPIs will have components with a shared cellular compartment by chance. We find that 482/990 (49%) of the PPIs in the “not found” category have proteins in a shared compartment. This analysis suggests that at least 28% of the PPIs in the “not found” category could be correct. Therefore, in the aggregate, together with a 5% FDR estimate established at the search level, we can estimate that 58-95% of the PPIs we detect in this set are true positives. The DSS-based crosslinking methods generate a similar range (**Supplementary Table 2**). We then investigated the quality of CSMs supporting the entries in the “not found” category and observed that 47-58% of the CSM scores were better than 0.5 (moderate to high quality, **Supplementary Figure 4**). Interestingly, click-linking scores are much superior to PhoX, although PhoX generates hits in the 76-95% range based on shared compartments (**Supplementary Table 2**). Finally, we performed a search of the click-linking data using CRIMP 2.0, a more conservative tool for PPI detection^31^. Fewer PPIs were detected than with pLink2 (863), but these also generated a sizable fraction in the “not found” category (**Supplementary Figure 5**). Taken together, the click-linking protocol likely identifies hundreds of novel, unmapped PPIs, generating data that are superior in quality and quantity to PhoX data even though affinity enrichment was not used.

Finally, we illustrate that click-linking does faithfully sample protein structure and capture interactors that reflect biological function. We tested the integrity of sampling by mapping the crosslinks to available protein structures using CLAUDIO^32^. Of the 14,528 crosslinked peptide pairs identified, CLAUDIO was able to map 7,802 across 1,465 protein structures, reflecting good structural sampling. Over 87% of these measured crosslinks mapped to within 35 Å (**Supplementary Figure 6**). As an example of a functional complex, we identified ACIN1, a nuclear protein that is crosslinked to SAP18 (a histone deacetylase complex subunit) and RNPS1 (RNA-binding protein with serine-rich domain 1) that together form the apoptosis and splicing-associated protein (ASAP) complex^33,34^. This sub-structure is part of a larger exon-junction complex (EJC) that regulates gene expression after transcription. Evidence for this is found in crosslinks between SAP18 and the Pinin splicing factor^35,36^. An AlphaFold Multimer model of the ternary complex supports the measured interprotein crosslinks, except for those situated in the highly disordered regions (**Supplementary Figure 7**).

In summary, click-linking is a very effective protocol for increasing the yield of protein crosslinks, and one that avoids the weaknesses of the traditional method. There are many reasons for the improved yield. In the first place, the method promotes greater control over the chemical reaction itself. Pre-stabilizing the proteome allows us to wash away partially hydrolyzed click precursors as well as labeled metabolites, both major classes of competing reaction products.

Additionally, self-competition is avoided^37^. That is, if two linkable residues are each labeled with a conventional homobifunctional crosslinker, they cannot crosslink. In click-linking, competition can only occur in the second reaction, but this will always generate a protein crosslink.

While the yields are clearly superior to other methods, it is somewhat surprising that PPI detection is not higher than the numbers we presented, given that our yields are significantly increased. The reasons for this are complex and the subject of a separate investigation, but we note here that search engines are not yet well optimized for searches of this scale. Nevertheless, combining cellular fixation with multistep reactions provides clear benefits over the classical crosslinking method, and illustrates a generic strategy that can be extended to many other reagent combinations in the pursuit of deep interactome sampling.

## Methods

### Cell culture

E. coli (DH5α) cells were seeded into 50 mL of 2YT + Amp100 (Fisher Scientific) medium and grown overnight at 37 °C, extensively washed with 1X PBS and lysed for the click optimization experiments. A549 cells were cultured in Ham’s F-12K medium (Gibco) supplemented with 10% fetal bovine serum (FBS, Gibco) and 1% penicillin-streptomycin (Gibco) at 37 °C in a humidified atmosphere with 5% CO_2_. For imaging experiments, cells were seeded in a 1cm microscope coverslip and grown to 80% confluency. For labeling and crosslinking experiments, they were seeded in 10cm Petri dishes and grown to the same confluency.

### Cell fixation and permeabilization

Cells had growth medium removed and were washed with 1X PBS buffer at pH 7.4. A 4% (v/v) formaldehyde solution in 1X PBS was freshly prepared and added to the cells for fixation. The incubation was held at 25°C for 10 min and excess formaldehyde solution was washed away with 1X PBS. Cells were then treated with a solution of 0.1% (v/v) Triton-X-100 in 1X PBS for 10 min at 25 °C. The surfactant was then washed out prior to labeling experiments.

### In situ monovalent labeling reactions

Fixed and permeabilized cells were washed three times with 1X PBS, then treated with both monovalent reagents simultaneously, at a final 1mM concentration for each. Stock solutions of both (2,5-dioxopyrrolidin-1-yl 3-(prop-2- ynyloxy)propanoate, Reagent **1**) and (2,5-dioxopyrrolidin-1-yl) 6-azidohexanoate, Reagent **2**) were prepared to 100 mM in anhydrous DMSO, then pre-mixed before adding to the plates.

Reactions were carried for 60 min at 25 °C under gentle mixing, before quenching with 20 mM ammonium bicarbonate. Excess reagents were washed out with 1X PBS and free azido groups were capped with 1 mM (11,12-didehydro-γ-oxodibenz[b,f]azocine-5(6H)-butanoic acid, DBCO-Acid). Cells were harvested from the plates in a lysis buffer composed of 100mM Tris(Hydroxymethyl)aminomethane, 150mM sodium chloride, 1% (v/v) Triton-X-100 and 2% (v/v) sodium dodecyl sulfate, at pH 7.8.

### Click optimizations

For the click optimization conditions, either E. coli or A549 cells were cultured as described above and the lysate was taken for labeling. 100µg of pre-reduced and alkylated lysate (see Sample work-up below) were bound to and cleaned-up following standard SP3-protein protocol. Instead of proceeding to the digestion step, the bead slurry was resuspended in 500µL of 1X PBS and proteins were treated with 1mM Reagent 1. Labeling reaction was carried for 60 min at 25 °C under strong mixing to keep the beads in suspension, followed by quenching with 20mM ammonium bicarbonate, mixing for another 10 min at 25 °C. Excess reagents were washed out from beads, which were resuspended again in 500µL of 1X PBS. Labeled proteins were then treated with 1mM 2-[2-(2-azidoethoxy)ethoxy]-ethanol with no click-catalysts present. For the time-course, copper sulfate, 3-(4-((Bis((1-(tert-butyl)-1H-1,2,3- triazol-4-yl)methyl)amino)methyl)-1H-1,2,3-triazol-1-yl)propane-1-sulfonic acid (BTTES) and sodium ascorbate were pre-mixed and introduced together to the solution at 5, 10 and 50mM, respectively. Reactions were quenched at the specific time points with 10mM DBCO-Acid. For the reagent ratio optimization, copper sulfate and sodium ascorbate were maintained at 5 and 50mM, respectively, while BTTES was tested in different concentrations. The click-chemistry reaction was conducted for 15min at 25 °C under strong mixing, before being quenched with 2mM DBCO-Acid. Excess reagents were washed out from beads and proteins were digested with trypsin as described below.

### In situ Click-Linking reactions

Cells were fixed, permeabilized and labeled with both reagents using the same conditions as above. After quenching the excess NHS esters and washes to remove excess reagent, the click-chemistry catalysts were added. Copper sulfate, BTTES and sodium ascorbate were pre-mixed and added together to the cells for final concentrations of 5, 10 and 50 mM, respectively. The clicking reaction was carried for 15 min at 25 °C under gentle mixing. Excess reagents and click-catalysts were washed away with 1X PBS, and 1 mM DBCO-Acid was added to cap any remaining free azido groups and prevent any downstream unwanted click reactions. Cells were then collected and lysed as described above.

### Sample work-up

All cell lysates were submitted to reduction of cysteines by incubation with 10 mM DTT at 52 °C for 30 min, followed by alkylation with 40 mM CAA at 25 °C for another 30 min in the dark. Reduced and alkylated lysate was bound to and cleaned up using the SP3-protein protocol. On-bead labeling experiments were performed at this point as described above, while samples where labeling and crosslinking occurred *in situ* proceeded directly to digestion. Digestion with trypsin in 50 mM ammonium bicarbonate, pH 7.8, at a 1:50 (w/w) trypsin-to-protein ratio was carried at 37 °C for 16h. A small aliquot of each sample was collected and desalted with C_18_ ZipTips (Millipore Sigma) and submitted to LC-MS/MS for the analysis of percent labeling and general quality control. The remaining samples were desalted using Pierce Peptide Desalting Columns (ThermoFisher Scientific) and submitted to fractionation.

### Fractionation

Samples were fractionated using a Vanquish Online 2D-LC system (ThermoFisher Scientific). For High-pH reverse phase fractionation, desalted peptide samples were resuspended in high pH Mobile Phase A (20 mM ammonium formate, pH 10) and loaded onto a ZORBAX RRHD Extended-C_18_ column (80Å pore size, 2.1 x 150 mm, 1.8 μm particles, Agilent). Separation was performed in a multistep 68 min gradient at 200 µL/min flow rate, from fixed 5% B (80% acetonitrile) for 1 min; 5-40% B for 42 min; 40-60% for 8 min; 60-95% for 2 min; fixed 95% for 4 min; then a ramp back 95-5% B for 2 min, holding at 5% for another 10 min. Eluate fractions were collected every minute from 1-61 min for a total 60 fractions, which were concatenated into 30 final fractions, dried down and resuspended in 0.1% formic acid for LC-MS/MS analysis. For the Clicked (SEC) sample, peptides were resuspended in SEC Mobile Phase (0.1% formic acid, 30% acetonitrile) and loaded onto a Superdex peptide PC 3.2/30 column (GE Healthcare). Separation was performed in 60 min using an isocratic flow of Mobile Phase at 0.08 mL/min. After 10 min, eluate fractions were collected every 1.875min for a total of 24 fractions. Out of the first 11 fractions, the first 7 were concatenated into 1, and fractions 8, 9, 10 and 11 were kept individually, while fractions 12 through 24 were discarded. Each of the 5 final fractions were submitted to the high pH fractionation and collection method described above. A total of 150 samples were dried and resuspended in 0.1% formic acid for LC-MS/MS analysis.

#### LC-MS/MS

Data were collected using a Vanquish Neo UHPLC coupled to an Orbitrap Ascend mass spectrometer fronted by an Easy-Spray Source (ThermoFisher Scientific). For the samples in Figure 4 and the 1xLabeling in Figure 2A, we also used an Orbitrap Astral mass spectrometer fronted by an Easy-Spray Source (ThermoFisher Scientific). For each sample, approximately 1 µg of peptide mass was injected onto a 300μm x 5mm PepMap Neo Trap Cartridge peptide trap column (C18, 5μm particle size, 100Å pore size, ThermoFisher Scientific), which were then submitted to reverse phase chromatography using an EASY-Spray HPLC analytical column (75μm x 50cm, C_18_, 2μm particle size, 100Å pore size, ThermoFisher Scientific) at a flow rate of 250 nL/min at 40 °C. Mobile Phase A consisted of 0.1% (v/v) formic acid in water, while Mobile Phase B consisted of 0.1% (v/v) formic acid in 80:20 acetonitrile:water (v/v).

For monovalent labeled samples, separation was carried using a multistep 120min gradient from 4-30% B for 85min; 30-45% B for 20min; then a 15min wash at 99% B. Mass spec data were acquired in positive mode using data-dependent acquisition (DDA) with a 2 second cycle time. Full MS scans were performed in the orbitrap set at 120,000 resolution (at *m/z* 200) for the 375-1875 m/z range, with maximum injection time set as Auto and normalized AGC target as Standard. Precursors with charge states 2-6+ were selectively isolated (quadrupole isolation window of 1.6 Th) for fragmentation using stepped HCD at Normalized Collision Energies (NCE) of 27, 30 and 33. Dynamic exclusion window was set to 10s. MS2 scans were performed using the orbitrap set at 30,000 resolution (*m/z* 200) with maximum injection time set to Auto and AGC target to 1e5. When the Astral was used, parameters were mostly the same, with the MS2 scan range set at m/z 150-2000, Normalized Collision Energy of 30%, maximum injection time of 10ms and AGC target of 300%.

For crosslinked samples (Clicked) coming from fractionation, separation was carried using a multistep 90min gradient from 4-20% B for 55min; 20-45% B for 15min; then a 20min wash at 99% B. Gradient was extended to 180min for the Clicked (SEC) samples. Mass spec data was acquired using DDA in a 2.5 second cycle time using roughly the same parameters, with few modifications. Full MS scans were performed in the orbitrap set at 120,000 resolution (at m/z 200) for the 375-1875 m/z range, with maximum injection time set to Auto and AGC target at 1e6. Precursors with charge states 4-8+ were selectively isolated (quadrupole isolation window of 1.6 Th) for fragmentation using stepped HCD at Normalized Collision Energies (NCE) of 27, 30 and 33. Dynamic exclusion window was set to 10s. MS2 scans were performed using the orbitrap set at 30,000 resolution (m/z 200) with maximum injection time set to Auto and AGC target to 1e5.

### Cell staining and imaging

A549 cells grown on glass coverslips were washed with 1X PBS then fixed and permeabilized as described above. Cells were treated with Reagent **1** and Reagent **2** (1mM each, added simultaneously) as described in the *in situ* monovalent labeling protocol, extensively washed with 1X PBS, then submitted to the click-chemistry reaction as described above, except for the labeled (but not clicked) controls. The clicked catalysts were extensively washed out with 5 mM EDTA in 1X PBS.

For phalloidin staining, samples were permeabilized with 0.5% Triton-X-100 for 10 min at 25 °C, then washed with 1X PBS. Cells were stained with CF647-phalloidin (1:40 v/v, Biotium) in PBS buffer for 60 min in the dark under gentle mixing. Excess phalloidin was removed and cells washed. Coverslips were then mounted onto microscope slides with EverBrite + DAPI mounting medium (Biotium) and sealed with clear nail polish and stored at 4 °C in the dark overnight. Cells were imaged using an AxioObserver.Z1 inverted microscope using the 40X oil immersion objective. Images were acquired in the 647 nm and DAPI channels with a 500 ms exposure time using the ZEN microscopy software. Images were pseudo-coloured, and brightness adjusted for merged composites using ImageJ software. The distribution of phalloidin fluorescent intensity was measured across 3 micrographs per condition using the ZEN blue software with a maximum bin count set to 4096 and bin size of 1 for the intensity histogram. The assessment of ultrastructure preservation or distortion came from monitoring select features from approximately 100 cells across 6-7 micrographs: major straited actin filaments, diffusive actin structures, presence of filopodia and the overall rounding of cell shapes.

For the tubulin immunofluorescent staining, samples were also submitted to a second permeabilization step with 0.5% Triton-X-100 in 1X PBS for 10 min at 25 °C, then washed. Samples were blocked with a 1% BSA (m/v) in 1X PBS solution for 30 min at room temperature.

The blocking solution was removed, and cells were stained with a 1:1000 dilution of anti-α- tubulin-FITC mouse monoclonal antibody (Sigma-Aldrich) in 1% BSA + PBS solution for 90 min at room temperature, then washed with 0.05% Tween-20 in 1X PBS. Coverslips were then mounted onto microscope slides with EverBrite + DAPI mounting medium (Biotium) and sealed with clear nail polish and stored at 4 °C in the dark overnight. Cells were imaged using the same microscope as above, using the 40X and 100X oil immersion objectives with 110 ms and 200 ms exposure times, respectively, in the FITC channel and 10 ms exposure time in the DAPI channel using the ZEN microscopy software. Image processing was done the same way as the Phalloidin staining.

### Metabolomics analysis

All analyses were performed at The Metabolomics Innovation Centre (Edmonton, Alberta, Canada) using standardized protocols. Briefly, metabolites were extracted (in quadruplicate) from fixed and permeabilized A549 cells, and from live A549 cells as a control, using methanol/water (4:1 v/v). Samples were normalized to 8 mM then labeled with light ^12^C_2_ dansylation reagent and each sample was combined with, and referenced against, to a pooled sample that was labeled with heavy ^13^C_2_ dansylation reagent for normalization purposes. Samples were run on a Bruker Impact II QTOF (positive ion mode, m/z range 220- 1000, 1 Hz acquisition rate) outfitted with a Thermo Scientific Vanquish LC using mobile phase A (0.1% formic acid in water) and mobile phase B (0.1% formic acid in acetonitrile). Separation was achieved using a gradient of B, starting at 25% (t = 0 min) and ending at 99% (t = 15 min), at a flow rate of 400 μl/min. Metabolites were identified by library match to the CIL library, the LI library and the MyCompoundID library, using accurate mass and retention time, generating 1814 identifications (or putative matches) from 2028 unique peak pairs.

### Monovalent Data Analysis

Monovalent labeling efficiency was accessed using MSFragger inside FragPipe (v22.0)^38^. For every sample, we performed an OpenMod search to investigate which labeling tags were being installed into proteins by the different reagents used and if there were any unexpected side products we did not anticipate. The OpenMod search was conducted against the full Escherichia coli proteome (UniProt database, retrieved September 2021) or the full Homo sapiens proteome (UniProt database, retrieved February 2023) with mostly default parameters. Notable changes include the MS1 range from -150 to 650Da, MS2 error tolerance 15ppm, inclusion of cysteine alkylation as a variable modification instead of fixed. After confirming the mass shifts, the LFQ-MBR workflow was used to investigate the quantitative proportion of labeled versus unlabeled peptides and thus estimating reaction performance according to the following equation:

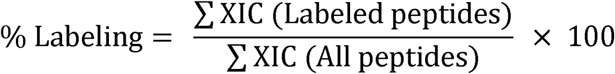

The LFQ-MBR search routine was also performed against the full proteome databases with the following parameters: cysteine carbamidomethylation (Δm = +57.021465 Da) as fixed modification; methionine oxidation (Δm = +15.9949 Da) and N-termini acetylation (Δm = +42.0106 Da) as variable modifications; 10 ppm for precursor mass tolerance; 15 ppm for fragment mass tolerance; trypsin as enzyme for protein digestion with 3 allowed missed cleavages. The mass shifts corresponding to the reagent modification were adjusted for every molecule as follows: Reagent **1** (Δm = 110.0366 Da), Reagent **2** (Δm = 139.0745 Da), DBCO- Capped-Azide (Δm = 444.1798 Da), Reagent 1 clicked with Azide-PEG2-Alcohol (Δm = 285.1324 Da). IonQuant was used for label-free quantitation and each mass spec run was processed individually.

### Crosslinking Data Analysis

After clicking, we monitored the residual non-clicked monolinks using the same pipeline described above through FragPipe. Crosslinking data analysis was performed using pLink 2.3.11^39^. Parameters used were as follows: peptide length 6-60; trypsin as the digestion enzyme with 3 allowed missed cleavages; carbamidomethylation of cysteines (Δm = +57.021 Da) as a fixed modification; oxidation of methionine (Δm = +15.995 Da) as a variable modification; 5 ppm for precursor and 10 ppm for fragment mass tolerance.

Data were searched against the full UniProt Human proteome databases at 5% false discovery rate (FDR) at the spectrum level. Crosslink (Δm = +249.111 Da) and monolink modifications were added for the clicked products, with specificity towards Lysine residues and protein N-termini (only one monolink is allowed on pLink, we selected the more abundant Reagent **1** at Δm = +110.037 Da).

The output from pLink was searched against the STRING human protein interaction database (v12.0, retrieved March 2024) using an in-house developed Python script and mapped into protein structures using CLAUDIO^32^, which is capable of mapping crosslinks both into known structures from the PDB^40^ and to AlphaFold generated models^41^ when a high-resolution experimental structure is not available.

The dataset from Clicked (SEC) sample was also searched against the full human proteome using CRIMP 2.0 inside Mass Spec Studio. Parameters used were mostly default with the exception of: peptide length 6-60; trypsin as the digestion enzyme; carbamidomethylation of cysteines (Δm = +57.021 Da) as a fixed modification; oxidation of methionine (Δm = +15.995 Da) as a variable modification; 5 ppm for precursor and 10 ppm for fragment mass tolerance, minimum fragment threshold = 3, maximum and minimum isolation window = 0.8, % E-Threshold = 50%.

### Cellular compartment co-localization analysis

Gene-ontology terms for proteins involved in PPIs were retrieved from the Uniprot database and the number of PPIs with one or more shared cellular compartment(s) was quantified using an in-house script, as a function of STRING score. To estimate how frequently shared compartments could be observed by chance, we pulled 1,736 proteins from the human proteome at random, then randomly assigned pairs to a PPI for compartment evaluation. This process was performed in triplicate and the results averaged.

### Determination of Reagent-accessible surface area

Reagent residue accessibility was determined using *freesasa*^42^, using publicly available structure predictions which were accessed via RCSB^43,44^. Predictions were solely from AlphaFold2 (version 4)^45^. Residue accessibility was calculated for all proteins whose structures were available. Peptides were aligned to the structure where required. The reagent accessibility was calculated using the Lee Richards algorithm for solvent accessibility, using a probe radius which was derived from the literature surface area of Succinimidyl Acetate (63.7Å^2^; probe-radius = 4.3Å). The result of this calculation is 0 when the measured entity (*e.g.* residue or atom) is completely occluded from the reagent in the static structure.

## Data Availability

The crosslinking and labeling data generated in this study have been deposited in the PRIDE partner repository^46^ with the dataset identifier PXD059118. The Source data for Figures 2A-D, 4, 5, 6A-D, S1, S2A-B, S4, S5A-B, S6 and S7C are provided in the Supplementary Information/Source Data file.

## Acknowledgements

This work was funded by the Natural Sciences and Engineering Research Council of Canada Discovery Grants RGPIN 2017-04879 to D.C.S. All figures created and/or assembled with BioRender.com, released under a Creative Commons Attribution-NonCommercial-NoDerivs 4.0 International license.

## Author Contributions Statement

D.C.S., B.C.A. and A.R.M.M. conceptualized the project and designed experiments. A.R.M.M. collected all imaging data. B.C.A. and A.R.M.M. collected all mass spectra data, with assistance from K.H.E. and analyzed all data. D.A.C. generated the scripts and performed the surface-accessibility analysis. D.C.S. and B.C.A. generated the first draft of the manuscript and all authors helped finalize the paper.

