## Supplementary information for "Click-linking: a cell-compatible protein crosslinking method based on click chemistry"

Dr. David Schriemer

Department of Biochemistry and Molecular Biology, Cumming School of Medicine

University of Calgary, 3330 Hospital Drive NW T2N-4N1

---

[#]Both authors contributed equally to the work.

**Supplementary information**

### Contents

**Supplementary Figure 1.** Parsing the surface accessibility of lysines labeled in a 1X precursor installation reaction.

**Supplementary Figure 2.** Optimization of the *in situ* click reaction on installed precursors.

**Supplementary Figure 3.** Metabolomics analysis of fixed and permeabilized cells using live cells as a control.

**Supplementary Figure 4.** Distribution of best CSMs associated with detected PPIs for each of three *in situ* crosslinking methods.

**Supplementary Figure 5.** Comparison of the PPIs detected using pLink 2 and CRIMP 2.0 (Mass Spec Studio) both at a 5% FDR at the highest level of organization.

**Supplementary Figure 6.** Distribution of crosslink distances from a large sample of structures and models.

**Supplementary Figure 7.** AlphaFold Multimer modeling of a ternary complex involving the ASAP complex and the splicing factor Pinin.

**Supplementary Table 1.** Distribution of reaction products for several *in situ* crosslinking reactions, based on numbers of unique identifications using pLink 2.3.11 at 5% FDR.

**Supplementary Table 2.** Analysis of protein co-compartmentalization for various *in situ* crosslinking strategies.

### Supplementary Figures

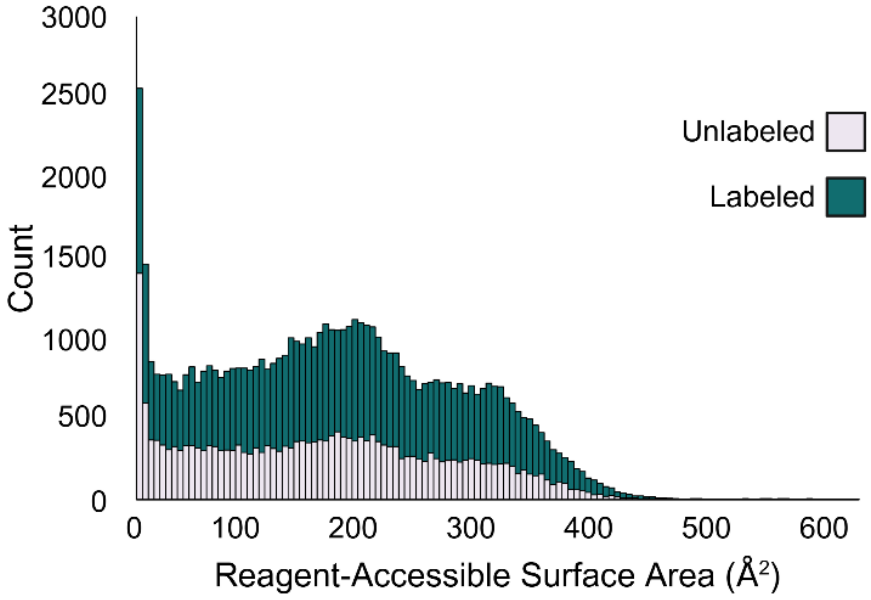


**Supplementary Figure 1. Parsing the surface accessibility of lysines labeled in a 1X precursor installation reaction.** Peptides were identified from the deep proteomic analysis of a 1x-labeled population of A549 cells and reduced to two sets of unique lysines (labeled and unlabeled). Using a probe based on succinimidyl acetate, reagent accessible surface area was measured for each lysine, where the x axis represents a scale in $Å^2$ relative to the reagent-inaccessible state. That is, all areas greater than zero are accessible, all areas equal to zero are inaccessible.

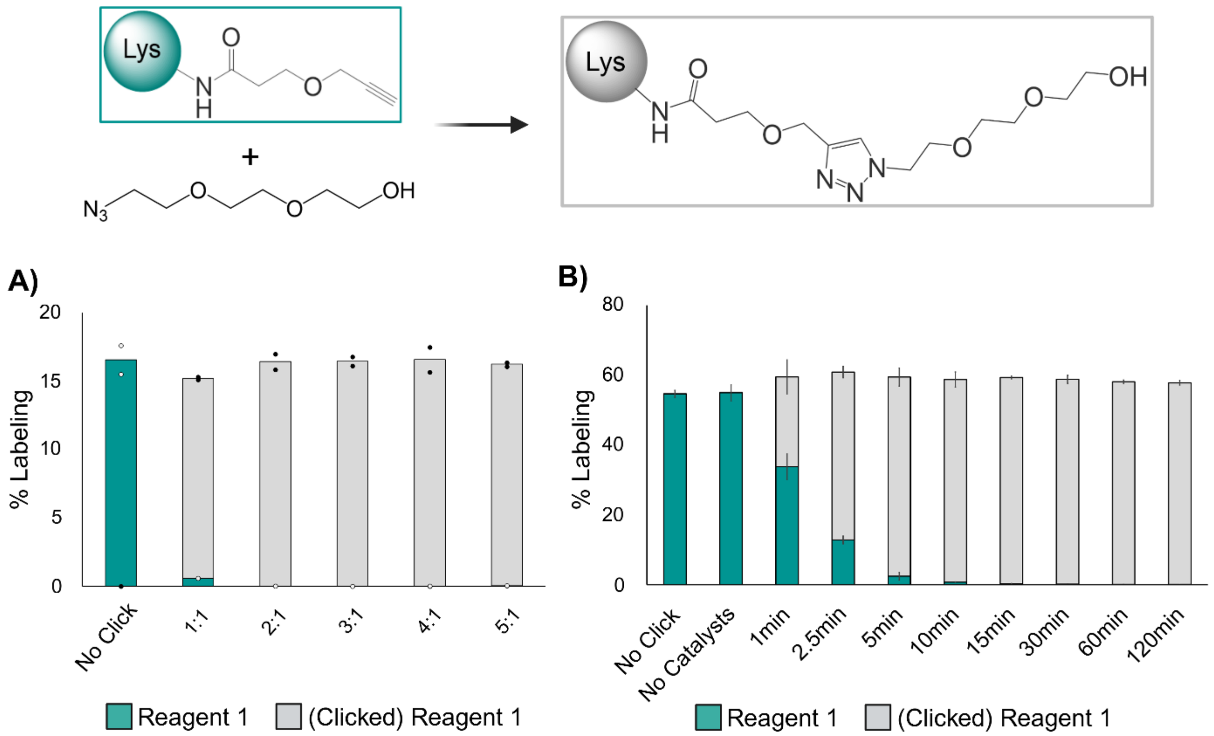


**Supplementary Figure 2. Optimization of the in situ click reaction on installed precursors.** E. coli (A) and A549 (B) full lysates were preinstalled with a propargyl reagent (Reagent 1), followed by the addition of ligand (BTTES) and Cu(I), together with 2-[2-(2-azidoethoxy)ethoxy]-ethanol as a capping agent (reaction product shown at top). (A) Varying the molar ratio of BTTES to Cu(I) and reacting for 15 min. Concentration of Cu(I) was fixed at 5mM, n = 2 biological replicates. (B) Varying the reaction time at a fixed 2:1 ratio of BTTES to Cu(I), n = 3 biological replicates. Reaction products measured by a deep bottom-up proteomics method.

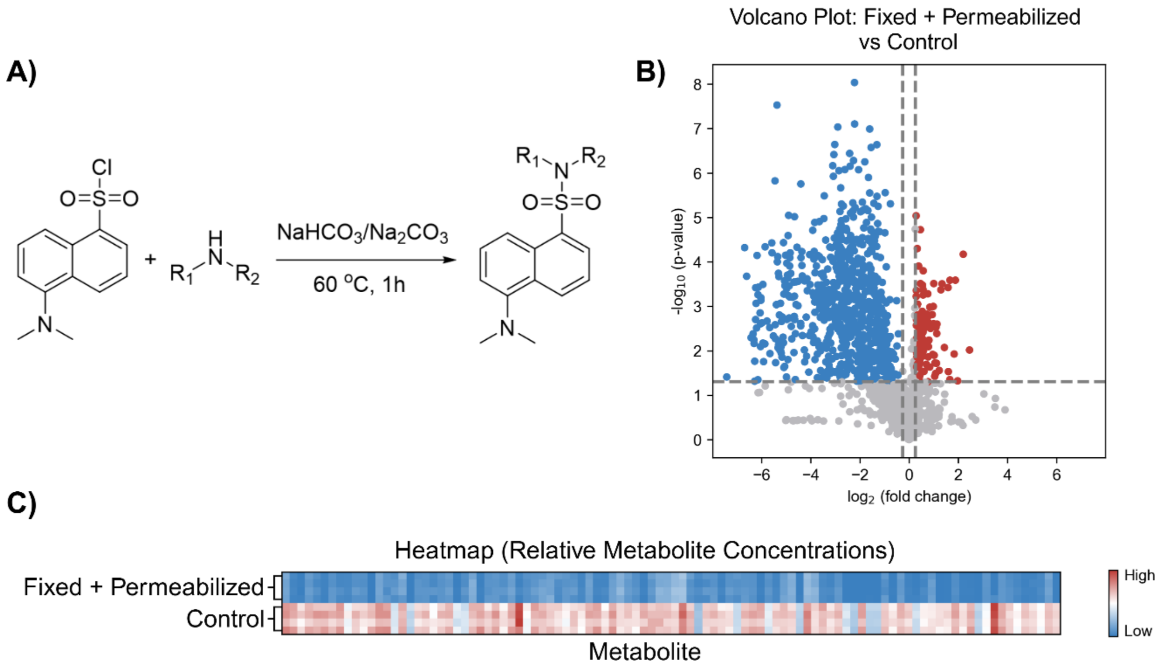


**Supplementary Figure 3. Metabolomics analysis of fixed and permeabilized cells using live cells as a control.** (A) Dansylation reaction scheme used to derivatize all amine-containing molecules, in both heavy and light form (see methods). (B) Volcano plot showing overall depletions (blue) in amine-reactive metabolites, and some enhancements (red). Enhancements are minor/artefactual, representing low abundance metabolites that tend to amplify the range in fold-change because of poorer ion statistics. (C) Heatmap of the top 100 high-intensity metabolites (same color scheme as B) showing overall depletion of amine-containing metabolites upon fixing and permeabilizing.

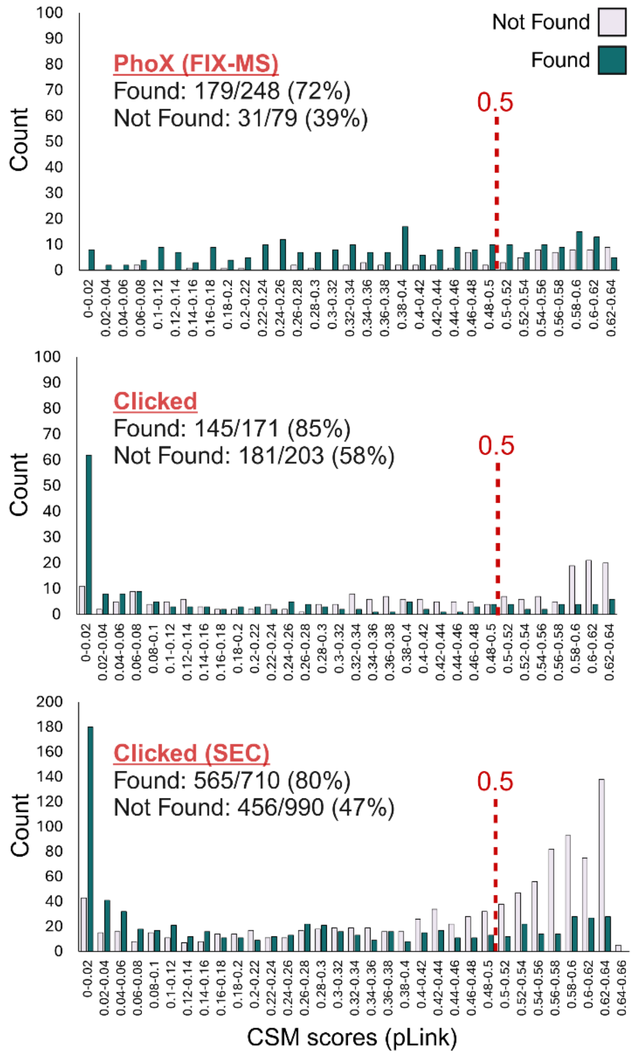


**Supplementary Figure 4. Distribution of best CSMs associated with detected PPIs for each of three *in situ* crosslinking methods.** Data shown as a function of PPIs in the Found category (STRING scores greater than 0.1) and Not Found category (no hits in the STRING database). A score cutoff of 0.5 is specified, to determine the fraction of CSMs with "quality" scores in each category.

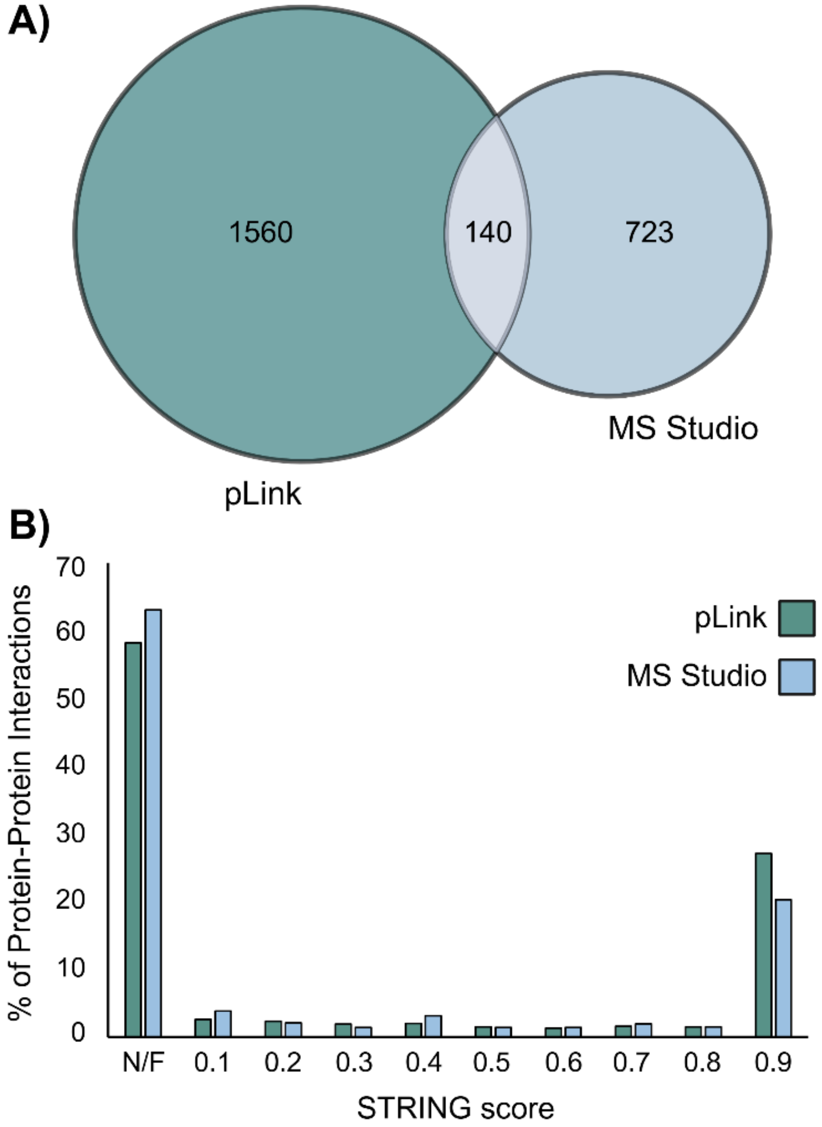


**Supplementary Figure 5. Comparison of the PPIs detected using pLink 2 and CRIMP 2.0 inside Mass Spec Studio, both at a 5% FDR at the highest level of organization**. (A) Venn diagram showing the number and overlap between search methods. (B) Distribution of PPI's detected in CRIMP 2.0 across STRING scores.

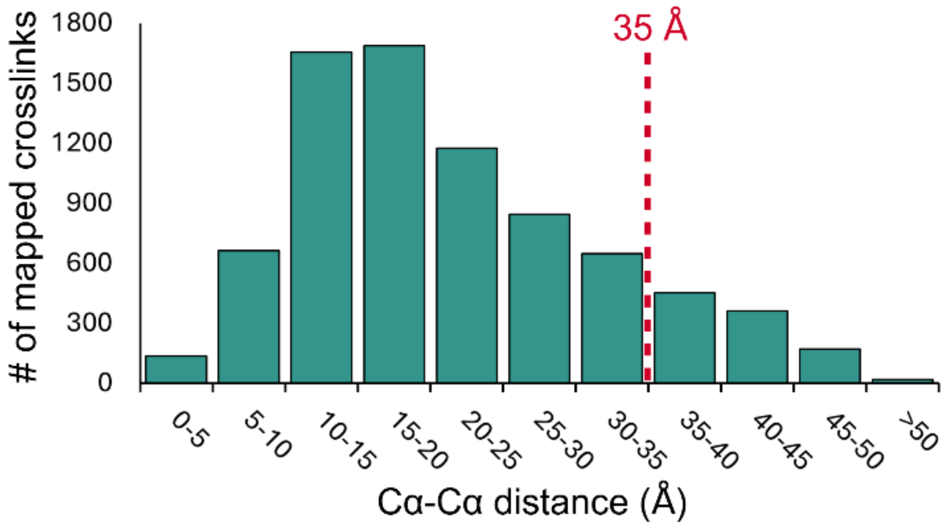


**Supplementary Figure 6. Distribution of crosslink distances from a large sample of structures and models**. Data derived from click-linked A549 cells processed by size-exclusion and high pH reverse phase chromatography prior to LC-MS/MS. 7,802 unique crosslinks were mapped across 1,465 structures from the PDB or, when experimental structures were not available, from AlphaFold-generated models, using CLAUDIO. Of all the mapped crosslinks, 6,802 (87.2%) were found within a distance of 35 Å.

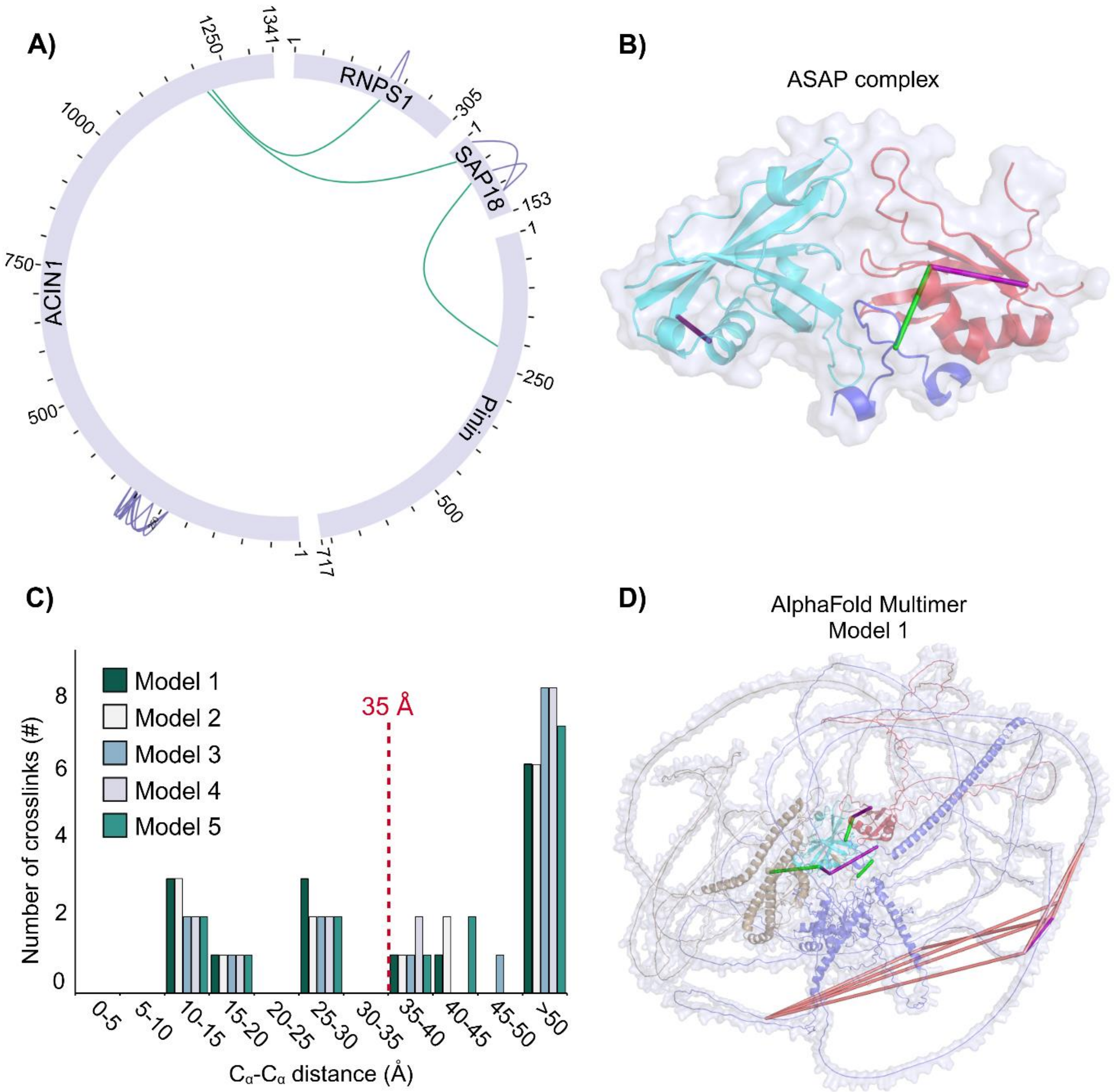


**Supplementary Figure 7. Alphafold Multimer modeling of a ternary complex involving the ASAP complex and the splicing factor Pinin.** (A) Circos plot of crosslink connectivity between SAP18, RNPS1, ACIN1 – constituents of the ASAP complex – and Pinin, and EJC-associated protein. (B) Mapping of the crosslinks into the crystal structure of the ASAP complex (PDB entry 4A8X), showing good agreement between the crosslinks and the high-resolution structure. Interlinks are in green, intralinks in purple. (C) Distance distribution of the experimental crosslinks when mapped into the top 5 models of the ASAP+Pinin tetramer complex generated by AlphaFold Multimer. (D) Mapping of the crosslinks into Model 1 from AlphaFold multimer, showing the core crosslinks in agreement with the model. Crosslinks associated with the structurally disordered regions violate the distance restraint, as expected.

**Supplementary information**

Interlinks are in green, intralinks in purple and overlength crosslinks in red. ACIN11 is in blue, SAP18 in cyan, Pinin in brown and RNPS1 in red.
